# GCN5a is a telomeric lysine acetyltransferase whose loss primes *Toxoplasma gondii* for latency

**DOI:** 10.1101/2025.01.14.633007

**Authors:** Vishakha Dey, Michael J. Holmes, Sandeep Srivastava, Emma H. Wilson, William J. Sullivan

## Abstract

*Toxoplasma gondii* is a protozoan parasite that causes persistent infection in warm-blooded vertebrates by undergoing differentiation from a replicative stage (tachyzoites) to a latent encysted stage (bradyzoites). Stage differentiation is critical for transmission and pathogenesis and relies on gene regulation driven by a network of transcription and epigenetic factors. We previously found in non-cystogenic type I RH strain parasites that the lysine acetyltransferase (KAT), GCN5a, is dispensable in tachyzoites but required to upregulate stress-response genes, suggesting a link with bradyzoite conversion. To address this possibility, we generated endogenously tagged GCN5a parasites and a genetic knockout in cystogenic type II Pru strain. We show that GCN5a protein, but not mRNA, increases during differentiation and complexes with unique protein partners, most of which contain AP2 domains. Pru strain tachyzoites lacking GCN5a augment bradyzoite-specific gene expression in the absence of stress. Loss of *GCN5a* slowed tachyzoite replication and heightened sensitivity to bradyzoite conversion but resulted in smaller cyst sizes compared to wildtype. Using CUT&Tag, we delineated the chromosomal occupancy of GCN5a relative to the essential KAT, GCN5b. While GCN5b localizes to coding regions, GCN5a surprisingly localizes exclusively to telomeres. These findings suggest that the loss of GCN5a leads to telomere dysfunction, which slows replication and promotes the transition to latency.

**IMPORTANCE:** *Toxoplasma gondii* is a single-celled parasite that persists in warm-blooded hosts, including humans, because it converts into latent tissue cysts. Switching from its replicating form into dormant cysts is a tightly regulated process that involves epigenetic factors such as lysine acetyltransferases GCN5a and GCN5b. This study is the first to examine the role of GCN5a in a cyst-forming *Toxoplasma* strain. We found that GCN5a protein, but not mRNA, increases during cyst development. Additionally, parasites lacking GCN5a replicate more slowly and are quicker to form cysts when stressed. We show that GCN5a and GCN5b work in different multi-protein complexes and localize to different areas of the genome; while GCN5b targets promoters of gene coding regions, GCN5a is exclusively found at telomeric regions. Our findings suggest a novel role for GCN5a in telomere biology that, when depleted, produces a fitness defect that favors development of latent stages.

## INTRODUCTION

*Toxoplasma gondii* is an obligate intracellular parasite that causes opportunistic infection. The life cycle is comprised of an asexual stage that occurs in the nucleated cells of warm-blooded animals and a sexual stage that takes place exclusively in felines (1). The sexual stage culminates in the release of oocysts that remain infectious for up to a year or more in the environment (2). Upon infection, tachyzoites reproduce asexually and disseminate throughout the host’s body. Tachyzoites can cross the placental barrier and infect the fetus, causing miscarriage or congenital birth defects (3). *Toxoplasma* evades host immune clearance by converting into latent bradyzoites that reside inside infected cells in the form of tissue cysts, establishing chronic toxoplasmosis (4). Tissue cysts are resistant to frontline drug treatments, allowing the parasite to persist within an infected host for life. If the host becomes immunocompromised, bradyzoite cysts can reactivate into proliferating tachyzoites that can cause life-threatening tissue destruction. Tissue cysts are another crucial means of parasite spread as naïve hosts can become infected after consuming contaminated meat (5). Given the importance of bradyzoites in parasite pathogenesis and transmission, understanding the molecular events coordinating stage switching remains an intense area of investigation.

The conversion of tachyzoites into bradyzoites requires alterations in gene expression, and several transcription factors mediate bradyzoite formation to varying degree. *Toxoplasma* contains 67 genes possessing a plant-like DNA-binding domain called Apetela-2 (AP2), several of which impact tachyzoite-bradyzoite conversion (4, 6). The master regulator of bradyzoite differentiation harbors a myb-like DNA-binding domain and is designated “bradyzoite-formation deficient 1” (*BFD1*). BFD1 binds to the promoters of stage-specific genes that are upregulated in bradyzoites, and parasites fail to initiate bradyzoite differentiation when *BFD1* is ablated (7).

Epigenetic machinery and histone modifications are also important for gene expression changes required for stage conversion, including histone acetylation mediated by general control non-repressed 5 (GCN5) proteins (4). Most early-branching eukaryotes possess a single GCN5 lysine acetyltransferase (KAT) that modulates stress responses (8, 9). In contrast, *Toxoplasma* contains two GCN5 KATs, designated GCN5a and GCN5b. To date, these GCN5 KATs have only been studied in type I RH parasites, which have largely lost their ability to readily convert to bradyzoites (10). While *GCN5b* is essential for parasite viability and regulates housekeeping genes, *GCN5a* is dispensable in tachyzoites (11, 12). Using a microarray for transcriptional profiling, RH parasites lacking GCN5a failed to upregulate ∼75% of the genes normally activated during alkaline stress (12).

As type I RH strain has largely lost its developmental capacity, it is not ideal for the study of factors involved in bradyzoite conversion. To address the potential role of GCN5a in stress-induced bradyzoite differentiation, we generated a tagged line and a genetic knockout in the cystogenic type II Prugniaud (Pru) strain. In contrast to RH strain, type II parasites lacking GCN5a displayed a reduction in replication and heightened sensitivity to stress-induced bradyzoite differentiation, although cyst sizes remained smaller than those made by wildtype. Transcriptional profiling of unstressed parasites revealed abnormal expression of bradyzoite genes in tachyzoites. We performed CUT&Tag (Cleavage Under Targets and Tagmentation) on both GCN5a and GCN5b, showing that GCN5a is exclusively associated with telomeric regions. Together, these findings suggest that loss of *GCN5a* likely leads to telomere dysfunction, thereby producing replicative delays and cellular stress that sensitizes the parasite towards bradyzoite conversion.

## RESULTS AND DISCUSSION

### GCN5a protein, but not mRNA, increases during bradyzoite differentiation

We previously generated a *GCN5a* knockout in type I RH *Toxoplasma*, finding that this lysine acetyltransferase (KAT) is dispensable for tachyzoite viability; however, parasites lacking GCN5a were impaired in their ability to upregulate 74% of the genes normally activated after 4 days in alkaline stress media, which is commonly used to induce tissue cyst formation in vitro (12). These results suggested that GCN5a is a stress-responsive KAT that may contribute to bradyzoite differentiation (13). Since type I parasites do not develop into mature tissue cysts at high frequency, we sought to characterize the role of GCN5a in a cystogenic type II strain. Using a CRIPSR/Cas9 approach, we first tagged endogenous *GCN5a* at its C-terminus with a 3xHA epitope in type II Prugniaud (Pru) parasites lacking KU80 and HXGPRT (Fig. S1A). GCN5a^HA^ was not detectable in unstressed tachyzoites but could be visualized by western blot and immunofluorescence assay (IFA) following alkaline stress for 24 h (Fig. S1B, C). IFA of stressed parasites showed that GCN5a^HA^ localizes exclusively to the parasite nucleus in the Pru strain (Fig. S1B). Western blotting of GCN5a^HA^ parasites revealed a band at 130 kDa, the expected size for the epitope-tagged protein (Fig. S1C). To ensure fidelity of the integration site, GCN5a^HA^ clones were further confirmed by PCR of genomic DNA (Fig. S1D).

Previous work on GCN5a in RH strain employed an ectopically expressed recombinant form of the protein, so endogenous expression patterns have yet to be evaluated (12). Therefore, we monitored mRNA and protein levels before and after a bradyzoite-inducing stress in our GCN5a^HA^ parasites. While *GCN5a^HA^* mRNA levels remain unchanged (Fig. 1A), GCN5a^HA^ protein expression is enhanced in response to alkaline stress (Fig. 1B). These results suggest that GCN5a protein is stabilized during alkaline stress, or its mRNA is subject to stress-induced preferential translation during bradyzoite conversion like *BFD1* and *BFD2/ROCY1* transcripts (7, 14–16). Consistent with this latter idea, the *GCN5a* mRNA contains a long and complex 5’-leader sequence that contains 26 upstream open reading frames (uORFs), features that have been linked to translational control during phosphorylation of eIF2α (17). In addition, a genome-wide RNA CLIPseq analysis identified BFD2-binding sites within the 5’-leader of *GCN5a* mRNA (16), which we’ve shown to be associated with cap-independent mechanisms of translational control for *BFD1* (15).

**Figure 1.**
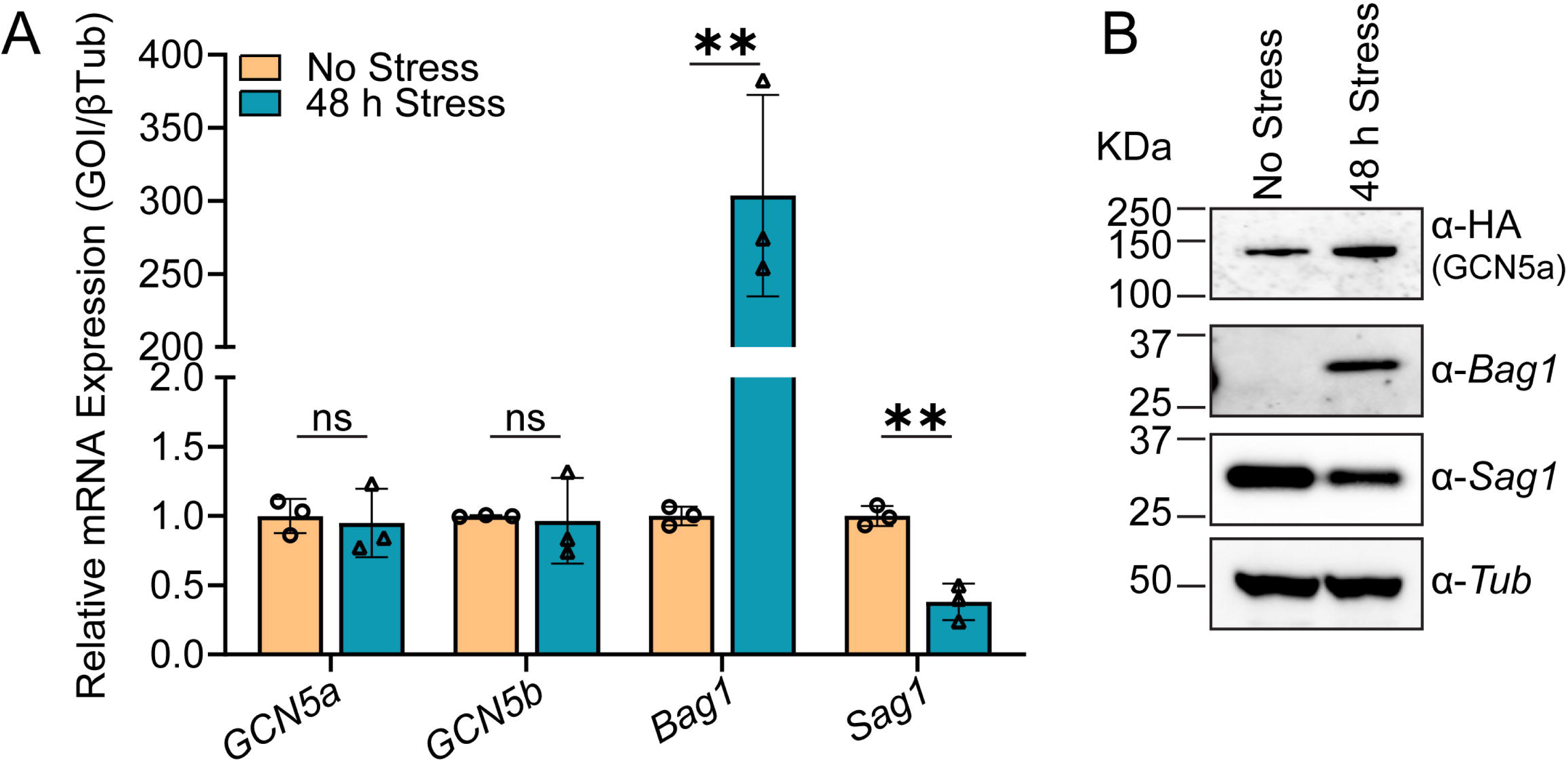
GCN5a^HA^ protein levels increase during alkaline stress. A. GCN5a^HA^ parasites were cultured in tachyzoite (No Stress) or alkaline stress conditions for 48 h (Stress). The level of *GCN5a* mRNA expression was monitored by RT-qPCR. The data are represented as the average of 3 biological replicates ± standard deviation. Statistical significance was measured by unpaired Student’s t test; ns = not significant; **p ≤ 0.01. B. Western blot of GCN5a^HA^ lysates from parasites cultured under designated conditions with stage-specific controls (BAG1 and SAG1) and a constitutively expressed control (TUB).

### Cystogenic parasites lacking GCN5a display reduced replication

We next generated a *GCN5a* knockout (ΔGCN5a) in the GCN5a^HA^ background. We replaced the entire GCN5a^HA^ coding sequence with an HXGPRT minigene cassette using a dual guide CRISPR/Cas9 strategy (Fig. S2A). The selected ΔGCN5a clone displayed no HA signal by IFA or western blot under stress conditions (Fig. S2B, C). The ΔGCN5a parasites were further validated by PCR of genomic DNA (Fig. S2D).

Previously, loss of GCN5a had no discernible effect on replication in type I RH tachyzoites (10). To determine whether the loss of GCN5a affected the viability of type II parasites, we conducted a plaque assay comparing GCN5a^HA^ and ΔGCN5a parasites under tachyzoite culture conditions. After 14 days, we found that ΔGCN5a parasites formed smaller plaques relative to GCN5a^HA^ parasites (Fig. 2A-B), although there was no difference in the number of plaques (Fig. 2C). These data suggest that the loss of GCN5a does not impact parasite invasion into host cells but does affect replication rate.

**Figure 2.**
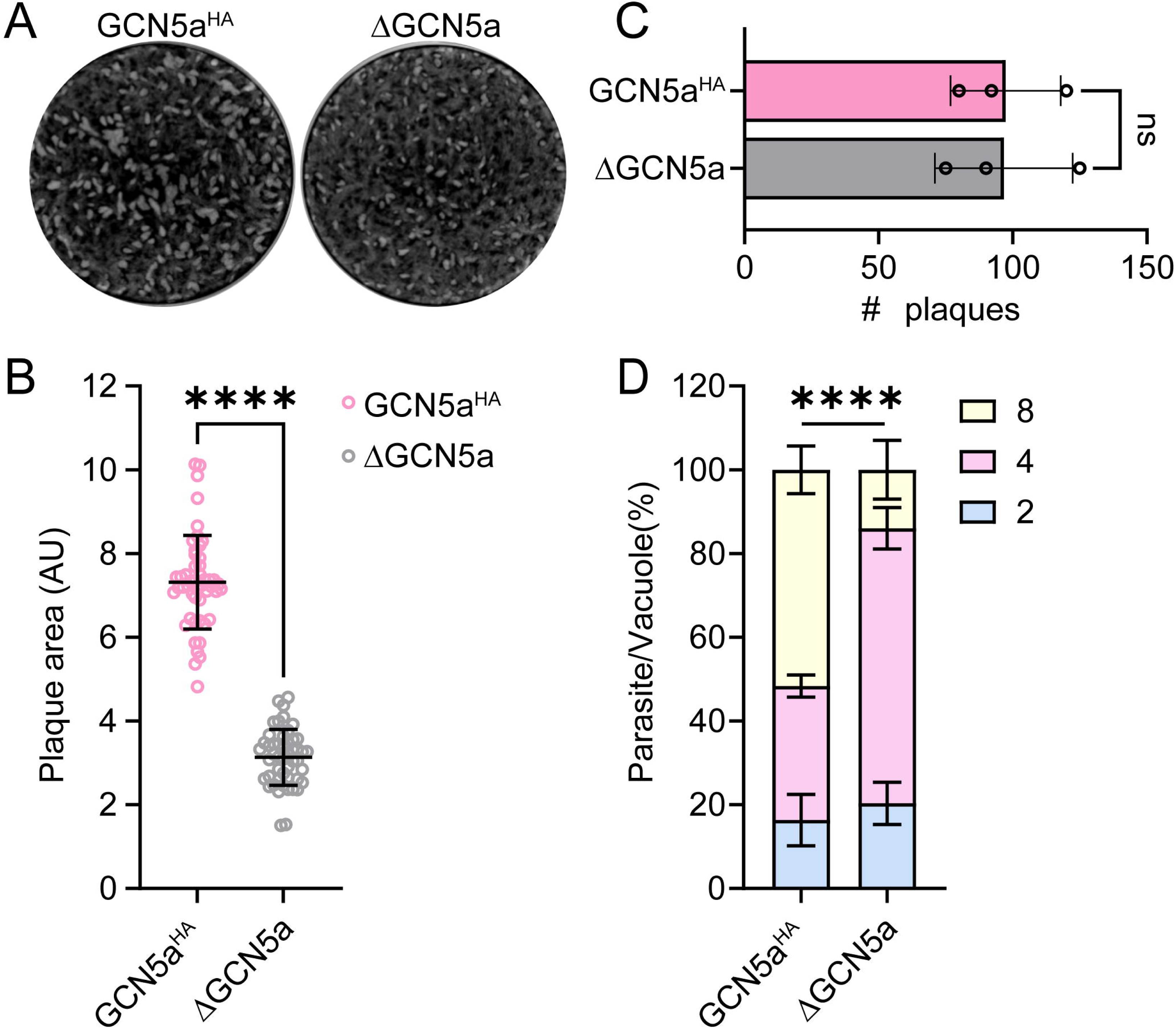
Phenotypic analysis of ΔGCN5a parasites. A. Plaque assay of GCN5a^HA^ and ΔGCN5a parasites grown under tachyzoite culture conditions for 14 days. B. The area of 50 randomly selected plaques was measured from 3 biological replicates. Statistical significance was measured by unpaired Student’s t test; **** represents p <0.0001. C. Number of plaques formed in HFF monolayers by GCN5a^HA^ and ΔGCN5a parasites. Each data point represents an average of 3 biological replicates ± standard deviation. Statistical significance was measured by unpaired Student’s t test; ns = not significant. D. Replication assay for GCN5a^HA^ and ΔGCN5a parasites. Number of parasites per vacuole was counted from 100 randomly selected vacuoles. Statistical significance was measured by one-way ANOVA with Tukey’s method for multiple comparisons; **** represents p <0.0001.

To examine this potential growth defect in ΔGCN5a parasites more closely, we performed a parasite replication assay. Parasites were allowed to grow for 16 hours and the number of parasites per vacuole was quantified. In contrast to the parental GCN5a^HA^ parasites, ΔGCN5a parasites contained more vacuoles with 2 parasites and fewer vacuoles with 4 parasites, confirming that the replication rate is slower for ΔGCN5a parasites (Fig. 2D). We further ensured that tagging of GCN5a did not affect the growth rate by comparing the PruΔKu80ΔHX parental strain with GCN5a^HA^ parasites using plaque and replication assays (Fig. S3). These results indicate that GCN5a contributes to parasite replication in type II cystogenic strains, unlike RH strain, suggesting strain-specific variations in the function of GCN5a.

### Increased expression of bradyzoite-associated genes in unstressed **Δ**GCN5a parasites

To further investigate the impaired replication phenotype observed for ΔGCN5a parasites, we performed differential expression transcriptional profiling of the parental GCN5a^HA^ (WT) and ΔGCN5a parasites (Table S2). Both upregulated a similar set of bradyzoite-specific genes in response to alkaline stress, including canonical markers *BAG1*, *ENO1*, and *LDH2* (Table S2; Fig 3A-B). However, the presence of GCN5a impacts the degree of the transcriptional response to stress. Although most genes were regulated in the same direction following bradyzoite induction, the degree to which these bradyzoite-associated genes are induced is blunted in the ΔGCN5a parasites compared to WT (Fig. 3B). For example, the log_2_ fold-change that occurs under stress for *ENO1* is >9 in WT but only ∼6 for ΔGCN5a parasites. The change in *BAG1* and *LDH2* expression is also much more pronounced in stressed GCN5a^HA^ parasites compared to the knockout, suggesting that the loss of GCN5a produces a muted stress response.

**Figure 3.**
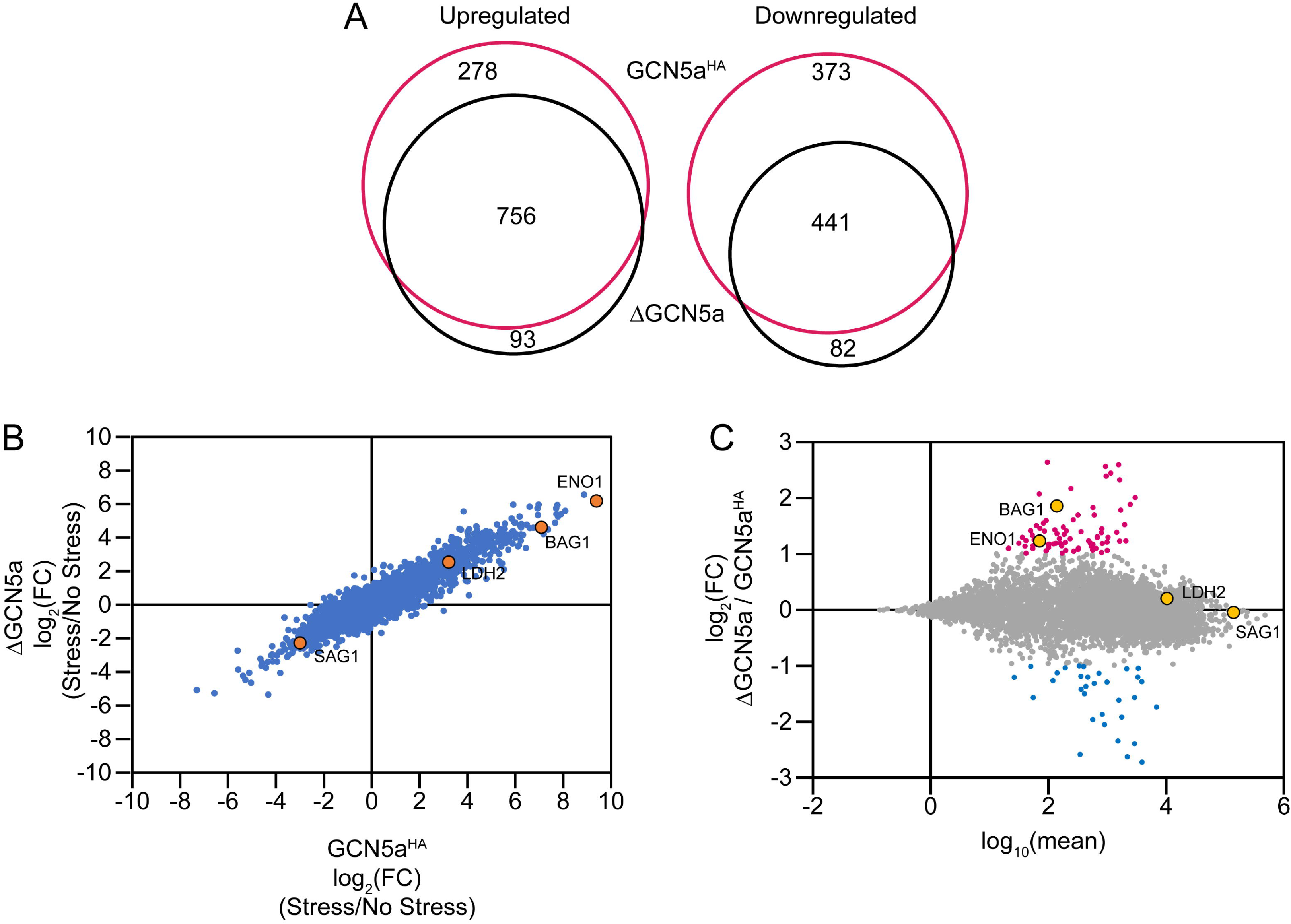
ΔGCN5a tachyzoites show increased expression of bradyzoite-associated genes. A. Venn diagram showing a high degree of gene co-regulation for GCN5a^HA^ and ΔGCN5a lines when they are subjected to bradyzoite-inducing culture conditions (alkaline media for 48 h). B. Comparison of differential gene expression between GCN5a^HA^ and ΔGCN5a parasites during stress-induced bradyzoite formation. Canonical stage-specific genes *BAG1*, *ENO1*, *LDH2*, and *SAG1* genes are highlighted in orange. C) MA plot depicting differential gene expression between GCN5a^HA^ and ΔGCN5a tachyzoites. Up- and downregulated genes are colored in red and blue, respectively. Canonical stage-specific genes *BAG1*, *ENO1*, *LDH2*, and *SAG1* genes are highlighted in orange.

Since a blunting of the stress response could either indicate a decrease in the induction of bradyzoite-associated genes or a change in basal gene expression in tachyzoites, we compared gene expression in GCN5a^HA^ and ΔGCN5a tachyzoites (Table S2; Fig 3C). Supporting the latter possibility, genes associated with early bradyzoite formation (18), such as *ENO1* and *BAG1*, were upregulated in unstressed ΔGCN5a tachyzoites. In contrast, genes that are turned on later during bradyzoite formation (18), such as *LDH2*, are unchanged in unstressed ΔGCN5a parasites. In fact, most (66 of 77) genes that displayed elevated basal expression in ΔGCN5a compared to WT tachyzoites were upregulated in both strains upon treatment with alkaline stress. Furthermore, elevated expression of early bradyzoite genes in the absence of stress is consistent with the slowed replication of ΔGCN5a parasites and supports the idea that ΔGCN5a tachyzoites may be poised for bradyzoite formation.

Counterintuitively, these findings are consistent with our previous observations in type I parasites lacking GCN5a. At the time, we interpreted a muted induction of bradyzoite-associated gene expression in response to alkaline treatment as a failure to initiate bradyzoite formation (12). Given our findings here, it is more likely that GCN5a deficiency in type I parasites also leads to increased expression of bradyzoite-associated genes in tachyzoites, blunting the response to alkaline stress, all while maintaining the rapid growth that characterizes of type I strains.

### Loss of GCN5a sensitizes tachyzoites towards bradyzoite conversion and results in reduced tissue cyst size

Bradyzoite conversion can be triggered through slowed parasite replication (19). In addition, our transcriptomic analysis revealed enhanced expression of several bradyzoite genes in ΔGCN5a tachyzoites, suggesting they may be predisposed towards differentiation. We infected HFF host cells with ΔGCN5a or WT (GCN5a^HA^) parasites and initiated bradyzoite differentiation using alkaline stress. The frequency of tissue cyst formation was monitored every 24 hours by staining for the cyst wall with *Dolichos biflorus* agglutinin (DBA). We observed that ΔGCN5a parasites differentiated more rapidly into cysts at 24 and 48 hours relative to WT (Fig. 4A left panel). By 72 hours, the differentiation rate of WT parasites had caught up to the ΔGCN5a parasites.

**Figure 4.**
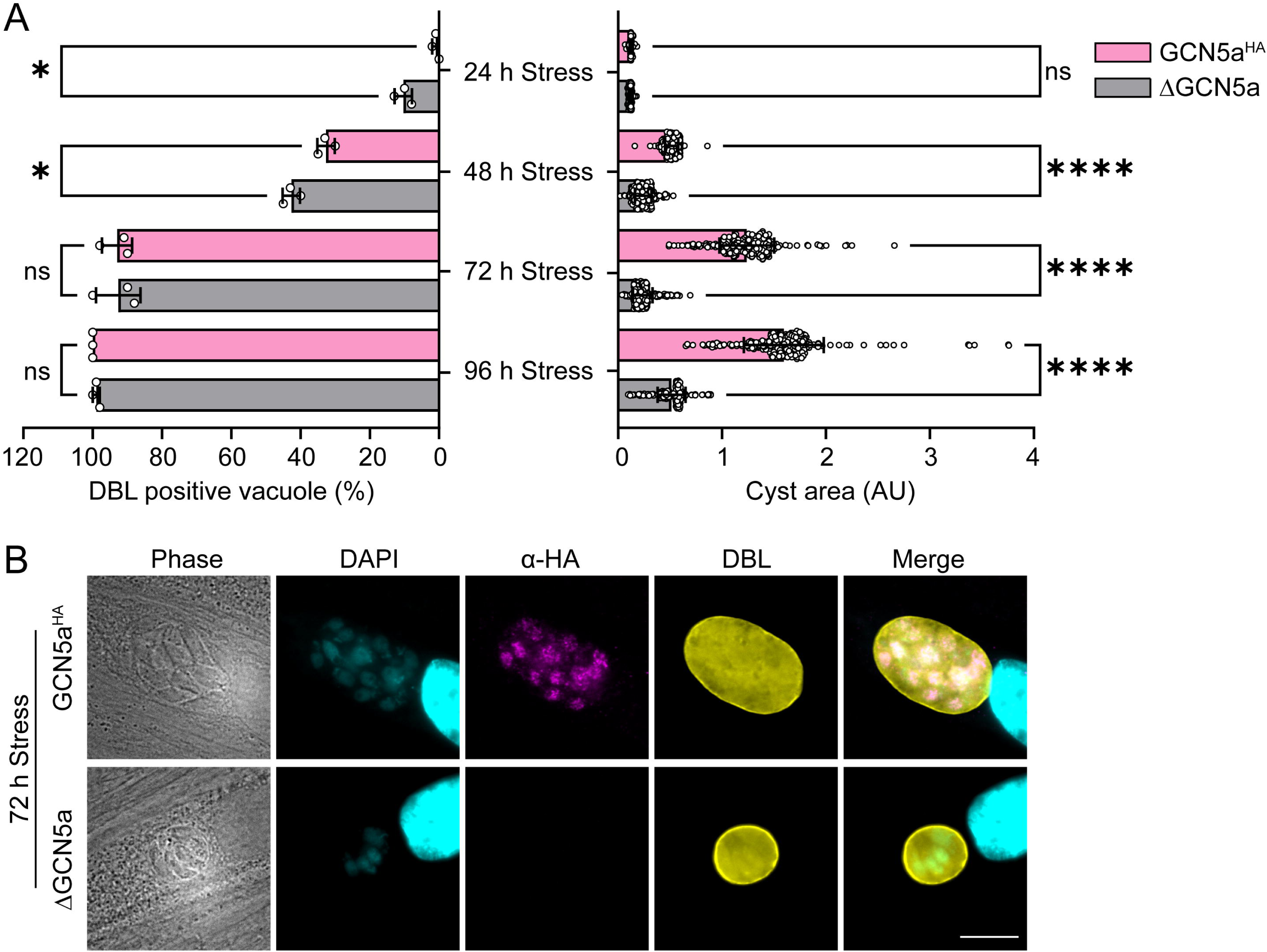
Cyst formation in ΔGCN5a parasites. A. (Left) Quantification of the number of DBA positive vacuoles following 24, 48, 72, and 96 h in alkaline stress. Each data point represents an average of 3 biological replicates ± standard deviation. Statistical significance was measured by one-way ANOVA with Tukey’s multiple comparison test; ns = p>0.05, * = p ≤0.05. (Right) The area of at least 100 randomly selected cysts was measured from 3 biological replicates. Statistical significance was measured by one-way ANOVA with Tukey’s multiple comparison test; ns = p>0.05, **** = p <0.0001. B. IFA of GCN5a^HA^ and ΔGCN5a parasite vacuoles after 72 h of alkaline stress. FITC-DBL was used to demarcate bradyzoite cyst wall (yellow). Anti-HA antibody was used to detect GCN5a^HA^ (magenta) and DAPI was used to stain nuclei (blue). Scale bar = 10 μm.

Quantification of average tissue cyst sizes revealed that those made by ΔGCN5a parasites were significantly smaller than those made by WT (Fig. 4A right panel). A representative IFA showing the smaller cyst sizes of ΔGCN5a parasites is shown in Figure 4B. These results confirm that the loss of GCN5a in a cystogenic strain makes tachyzoites more prone to differentiation and shows that the slowed replication produces smaller cyst sizes.

### Differences between GCN5a and GCN5b complexes

To shed further light on the function of GCN5a, we sought to elucidate its interacting proteins using co-immunoprecipitation (co-IP). As we could not obtain sufficient material using Pru GCN5a^HA^ parasites, we recreated this line in RHΔKu80ΔHX, which yields a higher biomass than Pru (Fig. S4A). We previously employed a similar strategy to characterize the GCN5b interactome (20). IFA confirmed that GCN5a^HA^ was localized to the nucleus as expected (Fig. S4B) and immunoblotting confirmed the presence of the endogenously tagged GCN5a^HA^ protein at the expected molecular weight of 130 kDa, increasing after stress as seen in Pru (Fig. S4C). Proper insertion of the tag at the *GCN5a* locus was further confirmed by PCR analysis of genomic DNA (Fig. S4D).

Results of three independent pull-downs of RH GCN5a^HA^ parasites under unstressed or alkaline stress conditions are shown in Table 1. Speaking to the fidelity of the pull-down data, we identified the coactivator protein ADA2-B (AP2VIIb-1), which we previously determined to interact with GCN5a and not GCN5b using a yeast two-hybrid assay (10). In contrast, GCN5b associates with exclusively with ADA2-A (10, 20).

**Table 1.** GCN5a interactome analysis. The number of peptides from three independent pull-downs is shown, generated from RH GCN5a^HA^ tachyzoites subjected to 48 h alkaline stress or not. Results are filtered for proteins represented by at least 5 peptides in two of the three replicates per condition. Proteins localized to organelles outside the nucleus were excluded. Grey highlight indicates an associating protein shared with the GCN5b complex.

| GeneID | Description (CRISPR fitness score) | No Stress | 48 h Stress |
| --- | --- | --- | --- |
| **TGGT1_254555** | **GCN5a (-1.0)** | **51\|26\|48** | **69\|50\|57** |
| TGGT1_234900 | PHD-finger domain-containing protein (-4.4) | 26\|15\|27 | 71\|65\|63 |
| TGGT1_262420 | AP2VIIb-1/ADA2-B (0.0) | 17\|14\|16 | 45\|56\|44 |
| TGGT1_247700 | AP2XII-4 (-5.0) | 7\|5\|11 | 41\|44\|36 |
| TGGT1_220520<br>TGGT1_220530 | AP2V-1 (0.8) | 1\|0\|5 | 21\|22\|15 |
| TGGT1_237090 | AP2X-5 (-3.5) | 0\|4\|0 | 14\|11\|9 |
| TGGT1_253440 | Cell cycle-associated protein kinase SRPK, putative (-3.0) | 5\|0\|3 | 9\|0\|11 |

We also found that GCN5a^HA^ pulls down with a PHD-finger domain-containing protein (TGGT1_234900), the putative cell cycle-associated SRPK (serine-arginine protein kinase), and several AP2 factors: AP2XII-4, AP2X-5, and AP2V-1. (The nanopore sequencing data available on ToxoDB (21) and nanopore sequencing data (22) indicate that TGGT1_220520 and TGGT1_220530 are misannotated as two genes and both should be attributed to AP2V-1). The PHD-finger protein and AP2XII-4 are also components of the GCN5b complex, but the others are unique to the GCN5a complex. Both GCN5 complexes contain a protein kinase, but they differ: GCN5a pulls down with SRPK whereas GCN5b associates with GSK (20, 23).

The associated components of the *Toxoplasma* GCN5a complex are novel and not conserved with other eukaryotic GCN5 complexes. The majority of the complex is composed of AP2 factors of unknown function. *Toxoplasma* possesses 67 AP2 domain-containing proteins, some of which have been linked to gene regulation while others to mitoribosome formation via RNA-binding activities (24). While ADA2-B is a homologue of the ADA2 coactivator known to enhance binding of acetyl-CoA to GCN5 for histone acetylation (25), the unique presence of an AP2 domain speaks to another possible function. The other AP2 factors contain no additional domains that provide clues to function with exception of AP2V-5. AP2V-5 harbors a WHIM1 domain, which has been implicated in chromatin remodeling as it pertains to nucleosome spacing (26). Notably absent from the GCN5a complex are proteins unequivocally associated with transcriptional activity, such as FACT (facilitates chromatin transcription) and TAF proteins that are found in the GCN5b complex (11, 20).

As one might expect, peptide counts increase following alkaline stress and the ensuing increase in GCN5a protein abundance. This represents a notable difference from the significant reorganization of the GCN5b complex during alkaline stress, which loses interactions with most of its AP2 factors, including AP2XII-4 (20). These findings are consistent with the model that the GCN5a complex is mobilized during stress (12) and support that GCN5a and GCN5b evolved to carry out distinct functions in the parasite.

### Chromosomal occupancy of GCN5a and GCN5b

Given the differences in complex composition and phenotype between the two GCN5s, we performed a comparative analysis of genome occupancy for each under stressed and unstressed conditions. To make a line comparable to Pru GCN5a^HA^, we engineered PruΔKu80ΔHX parasites to express endogenous GCN5b tagged with HA at its C-terminus (Fig. S5A). Immunoblotting confirmed the presence of the tagged GCN5b^HA^ protein at the expected molecular weight of 115 kDa (Fig. S5B). IFA demonstrated the expression of GCN5b^HA^ in the nucleus as expected (Fig. S5C). The accurate tagging of the *GCN5b* gene was further validated through PCR analysis (Fig. S5D). In contrast to GCN5a, we noted that protein levels for GCN5b do not change in response to alkaline stress (Fig. S5E).

CUT&Tag profiling revealed that GCN5b associates with the transcriptional start sites of protein coding genes but GCN5a does not (Fig. 5A). Surprisingly, GCN5a is robustly enriched at chromosomal ends over the repetitive sequence characteristic of telomeres (Fig. 5B). Moreover, the association of GCN5a with telomeres becomes more pronounced in the 48 h stress samples when GCN5a protein levels increase relative to the unstressed conditions. We did not detect GCN5b at telomeric regions in either culture condition.

**Figure 5.**
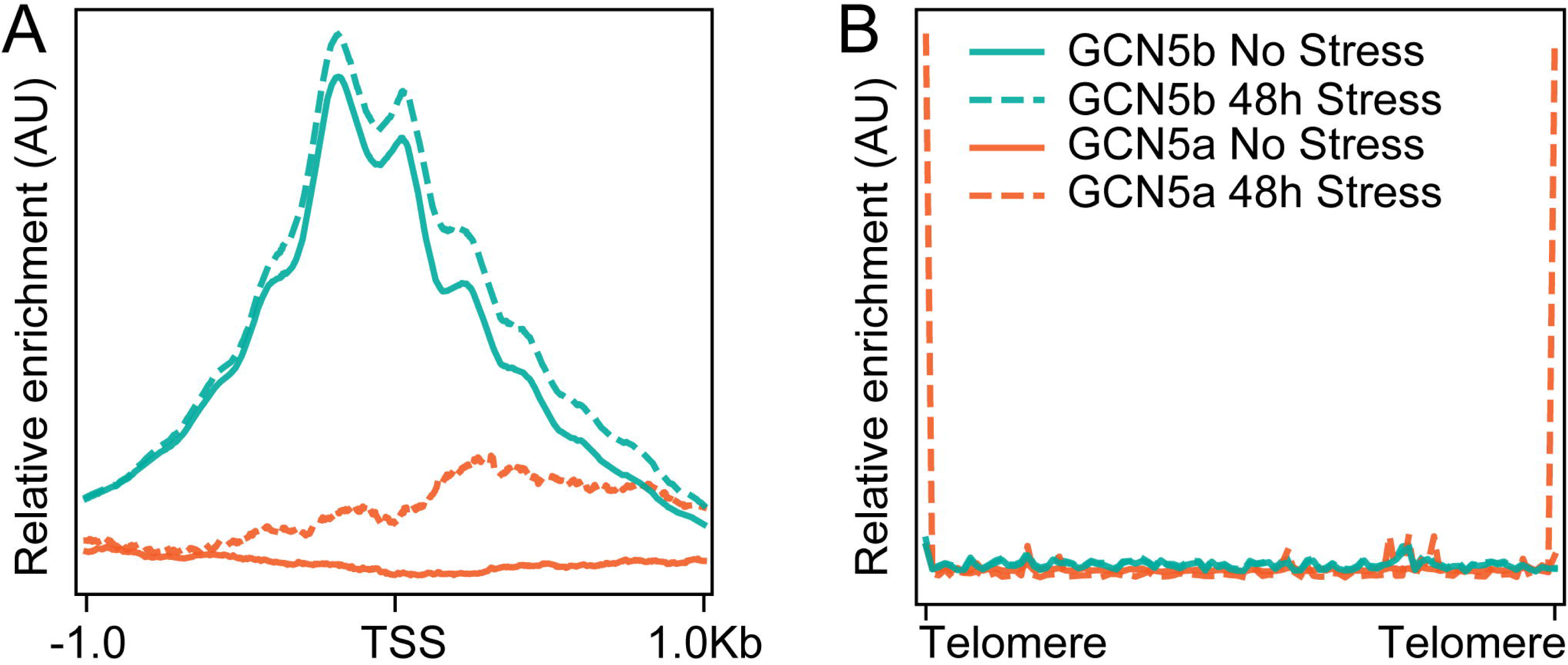
Chromosomal occupancy of GCN5a versus GCN5b. A. Meta-gene plot demonstrating enrichment of GCN5b, but not GCN5a, near the transcriptional start sites (TSS) of protein coding genes as determined by CUT&Tag. Legend depicting sample identification is shown in panel B. B. Meta-chromosomal plot demonstrating enrichment of GCN5a, but not GCN5b, at telomeres.

To confirm the association between GCN5a and telomeres, we performed fluorescent in situ hybridization (FISH) using a telomeric probe in GCN5a^HA^ parasites (Fig. 6A). Co-localization analysis revealed partial overlap between GCN5a^HA^ and telomere signals, consistent with the CUT&Tag data (Fig. 6B). Five additional nuclei were examined, yielding similar results (Fig. S6). To further confirm this interaction, we conducted an independent ChIP of GCN5a^HA^ (and parental parasites as a negative control) followed by PCR using primers that amplify the genomic segment spanning the subtelomeric region of chromosome ChrIb into its telomere. In agreement with the CUT&Tag data, the ChrIb telomeric region was detected by PCR following ChIP of GCN5a^HA^ (Fig. 6C). Also consistent with CUT&Tag data, we could not amplify *tubulin* or *B1* gene sequences following ChIP of GCN5a^HA^, supporting that GCN5a localizes specifically to telomeric regions and not coding sequences. These experiments collectively demonstrate that GCN5a localizes to telomeric regions and affirms GCN5b operates at coding sequences.

**Figure 6.**
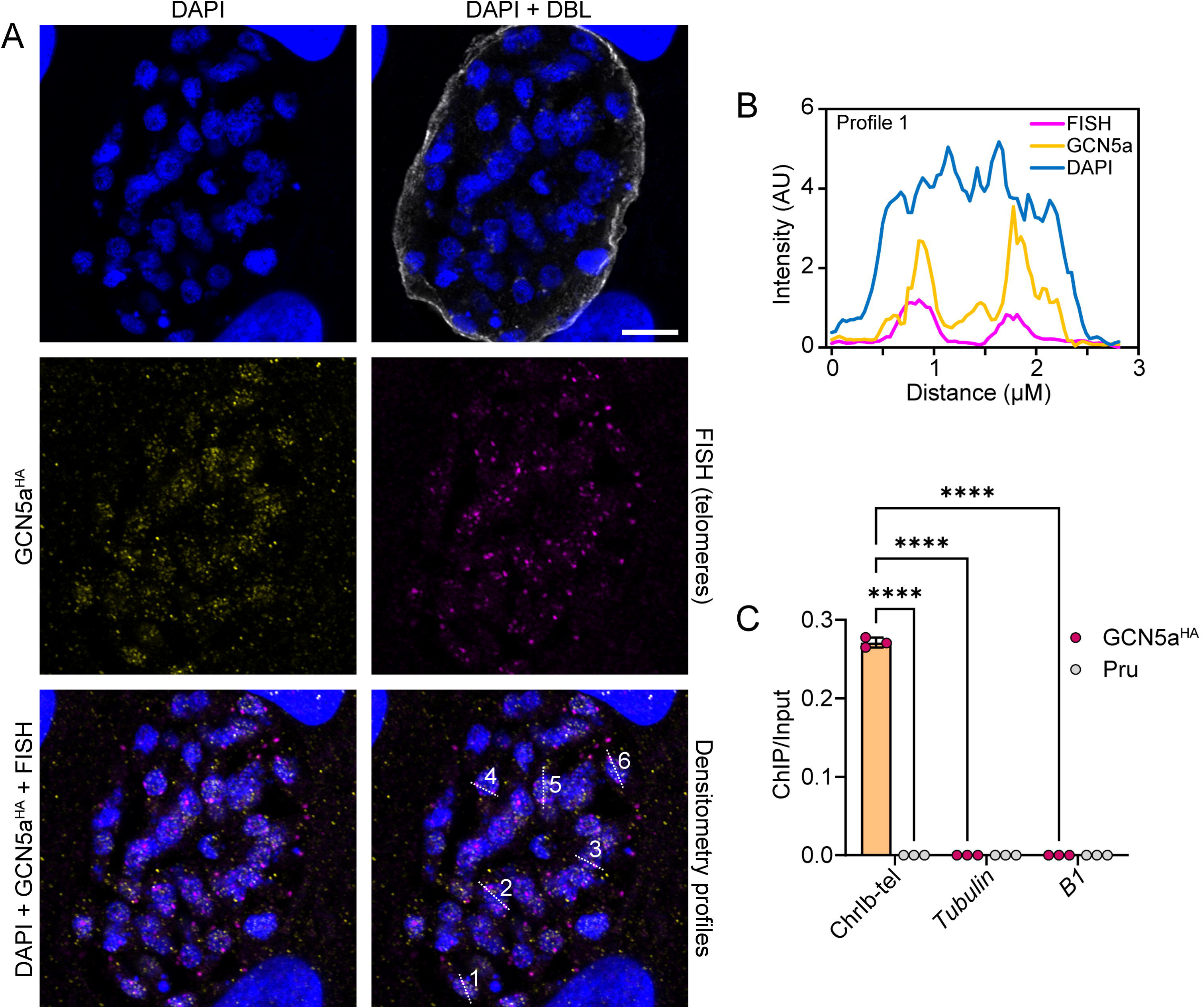
Independent validation of GCN5a localization to telomeres. A. GCN5a^HA^ parasites were cultured under alkaline stress conditions for 72 hours before IFA analyses. A representative vacuole of telomere FISH is shown. DNA is stained with DAPI (blue), GCN5a^HA^ is stained with anti-HA (yellow), telomere is stained with Cy5-telomere probe (magenta). Scale bar = 10 μm. The lower right panel indicates the profile regions that were assessed for signal intensity. B. Densitometry profile indicated in lower right panel of A, showing overlap between GCN5a and telomeric fluorescent signals. Profiles 2-5 are in Supplemental Figure S6. C. GCN5a^HA^ and PruΔKu80ΔHX (Pru) parasites were cultured alkaline stress conditions for 48 h prior to harvesting for ChIP-qPCR using anti-HA antibody. Immunoprecipitated DNA was examined by PCR using primers spanning the telomeric region of ChrIb (ChrIb-tel) or designated controls. Results are represented as a ratio of ChIP/Input DNA. Each data point represents an average of 3 biological replicates ± standard deviation. Statistical significance was measured by one-way ANOVA with Tukey’s multiple comparison test; ns = p>0.05, **** = p <0.0001.

### Concluding remarks

*Toxoplasma* is unusual among early-branching eukaryotes and fellow apicomplexan parasites in possessing two GCN5 KATs. We previously postulated that the non-essential KAT GCN5a resulted from a gene duplication of the essential KAT GCN5b (10). The present study shows that each GCN5 family member operates in a separate multi-protein complex to carry out distinct cellular functions. GCN5b fits the profile of a standard eukaryotic KAT, partnering with proteins directly linked to transcriptional activity (11, 20) and localizing to gene coding regions. Aside from AP2XII-4 and the PHD-finger protein TGGT1_234900, GCN5a partners with unique proteins and resides at telomeric regions. In addition, GCN5b protein levels do not increase during stress whereas GCN5a protein levels do.

A model fitting our findings as well as those from previous studies suggests that GCN5a evolved to promote telomere integrity, hence its loss generates a stress that slows replication and primes the parasite for latency as a survival strategy. Evidence from other species lends support to this model. In mammals, GCN5 interacts with proteins at telomeric regions and its ablation in mouse or human cells leads to telomere dysfunction (27, 28). Lysine acetylation is a means by which human telomeric protein TRF2 is stabilized (29). The GCN5-family KAT PCAF facilitates the resolution of R-loops that can accumulate at telomeres (30). Our findings open a new avenue for continued interrogation of *Toxoplasma* GCN5a and how it may participate in telomere biology.

## MATERIALS AND METHODS

### Host cell and parasite culture

*Toxoplasma gondii* PruΔKu80ΔHX (31) and RHΔKu80ΔHX (32) strains parasites were cultured in confluent human foreskin fibroblast (HFF) cells (ATCC, SCRC-1041) containing Dulbecco’s modified Eagle medium (DMEM) (GIBCO, 10-017-CM) supplemented with 5% inactivated fetal bovine serum (FBS) (R&D systems, S11150). Parasite cultures were maintained in a humidified 5% CO_2_ incubator at 37°C. HFF cells were grown in DMEM supplemented with 10% inactivated FBS. Alkaline medium (pH 8.2) for bradyzoite culture conditions contained RPMI 1640 (GIBCO, 31800-022) supplemented with 5% FBS and 50 mM HEPES. Parasites in alkaline medium were grown in ambient CO_2_ and the medium was changed daily.

### Transgenic parasite generation

Primers for this study were ordered from Integrated DNA Technologies (Table S1). GCN5a (TGME49_254555) and GCN5b (TGME49_243440) were endogenously tagged with a triple hemagglutinin (3xHA) epitope tag at their C-termini using CRISPR/Cas9 and double homologous recombination. A single guide RNA (sgRNA) sequence near the stop codon was chosen using the EuPaGDT tool (33) and the sequence was cloned into pCas9-GFP plasmid (34), designated as pCas9-GCN5a/b. A donor amplicon with homologous regions to upstream of the stop codon and 3’-UTR of the gene was PCR amplified to incorporate 3xHA sequence and the DHFR*-TS selectable marker (35).The Cas9 plasmid and the donor amplicon were co-transfected using a Lonza Biosciences Nucleofector and cultured for three passages in medium containing 1.0 μM pyrimethamine (Sigma, SML3579) and cloned by limiting dilution. The modified locus of the transgenic clone was validated by PCR analyses. The resulting parasite lines generated in Pru are labeled GCN5a^HA^ and GCN5b^HA^, while the line generated in RH is denoted as RH-GCN5a^HA^.

To generate the knockout GCN5a, two gRNAs were designed to replace the entire *GCN5a* coding sequence and DHFR*-TS cassette in GCN5a^HA^ parasites using a strategy that has been outlined previously (36). Both gRNAs were cloned into pCas9-GFP plasmid (34) to target immediately upstream of the *GCN5a* coding sequence (pCas9-gRNA1) and downstream of the *DHFR*-TS* 3’-UTR (pCas9-gRNA2). The resulting plasmid was denoted pCas9-dual-Guide. A double-stranded donor containing an HXGPRT selectable marker (37) flanked by homologous regions to the 5’- and 3’-UTRs of *GCN5a* was PCR amplified. Parasites were co-transfected with pCas9-dual-Guide and the repair template in GCN5a^HA^ parasites and selected for three passages in medium containing 25 μg/ml mycophenolic acid (Sigma, 475913) and 50 μg/ml xanthine (Sigma, X7375) and cloned by limiting dilution. The resulting parasite line was validated with genomic PCRs and is termed ΔGCN5a.

### Parasite growth assays

Standard growth assays were conducted as previously described (38). Briefly, for plaque assays, 500 parasites were used to infect confluent HFF cells grown on 12-well plates in tachyzoite medium. Plates were incubated undisturbed for 14 days until plaques were formed, which were visualized by crystal violet staining. Plaques were counted and the area was measured by ImageJ.

For replication assays, parasites were allowed to invade confluent HFFs for 4 h in tachyzoite medium. After invasion, the medium was replaced to wash away extracellular parasites. The number of parasites per vacuole was counted under light microscopy at the indicated time points.

### Immunofluorescence assay (IFA)

Parasites were allowed to infect confluent HFF cell monolayers on coverslips, cultured under tachyzoite or bradyzoite conditions with media changes every 24 h. Coverslips were fixed with 4% paraformaldehyde (Sigma, P6148), washed with PBS, and treated with blocking buffer (3% BSA, Sigma, A9418) and 0.2% Triton X-100 (Sigma, 93443) for permeabilization. Primary antibody incubation (1:1,000 rat anti-HA, Roche, 11867423001) was performed at 4°C for 16 h. After washing, cells were incubated with secondary antibody (1:5,000 Alexa Fluor 598 anti-rat, Thermo Fisher, A-11007) and 1 μg/ml DAPI (Invitrogen, D1306) for 1 h in the dark. For cyst walls, 1:500 FITC-conjugated *Dolichos biflorus* lectin (Vector Laboratories, FL1031) was added. After washing, coverslips were mounted with ProLong Gold Antifade (Invitrogen, P36930) and left to set overnight before imaging with a Nikon Eclipse E100080i microscope.

### Western blotting

Parasites were lysed with RIPA buffer supplemented with protease and phosphatase inhibitor cocktail (Thermo Fisher, 78425). The lysate was sonicated, centrifuged at 13,000×g for 2 min, and the supernatant was boiled for 10 min with 1X SDS-PAGE loading buffer and β-mercaptoethanol (Sigma, M3148). Samples were separated on NUPAGE 4-12% Bis-Tris gels (Invitrogen, NP0326BOX) and transferred to nitrocellulose membrane (Cytiva, 10600001). Blots were blocked in 5% non-fat milk/TBST for 1 h at room temperature, incubated with primary antibody at 4°C for 16 h using 1:1,000 dilution rat anti-HA (Roche, 11867423001) for HA-tagged proteins and 1:1,000 rabbit anti-BAG1 polyclonal antibodies (39) (provided by Vern Carruthers, University of Michigan) or 1:5,000 rabbit anti-tubulin polyclonal antibodies (provided by David Sibley, Washington University). After washing the blot with TBST, secondary antibody was applied using the appropriate HRP-conjugated antibodies (GE healthcare) at a 1:5,000 dilution. Blots were imaged using SuperSignal West Femto (Thermo Fisher, 34095) and detected via Bio-Rad ChemiDoc Imaging System.

### Real-time reverse transcription PCR (qRT-PCR)

Confluent HFF cells in T-25 flasks were infected with 10^5^ parasites. After 4 h, the medium was replaced with tachyzoite or bradyzoite medium. Total RNA was isolated using TRIzol-chloroform (Invitrogen, 10296028). cDNA synthesis was performed with SuperScript IV First-Strand cDNA synthesis system (Invitrogen, 18091300). cDNA was diluted 1:100 and used for real-time PCR with SYBR Green PCR Master Mix (Applied Biosystems, 4368708) on an Applied Biosystems QuantStudio 5 system. β-tubulin (TGME49_266960) was used for normalization. We verified that β-tubulin mRNA expression did not change under alkaline stress when either actin (TGME49_209030) or GAPDH (TGME49_289690) was used for normalization. Primers are listed in Table S1.

### Stage-specific RNAseq

An equal number of parasites (10^6^) were used to infect a confluent HFF cell monolayers in T-75 flasks for 4 h. The media were then changed to tachyzoite or bradyzoite medium and cultured for 48 h. Parasites were harvested by syringe passaging followed by filtration. Total RNA was isolated using TRIzol-chloroform and polyA-enriched RNAseq libraries were generated by Azenta Life Sciences using standard Illumina protocols. Libraries were sequenced 2×150 bp. Sequencing reads were trimmed of adapter sequences with Cutadapt (40) and mapped to the *Toxoplasma* genome (v55) (21) with HISAT2 (41). Differential gene expression was assessed with DESeq2 (42) after obtaining read counts with HTseq-count (43). Data pertaining to this experiment can be obtained at GSE286090 (reviewer token: mtgzmkkopxqtpyn).

### Immunoprecipitation of HA-tagged proteins

RH-GCN5a^HA^ and untagged RH parasites were cultured under both no stress and 48 hr alkaline stress conditions. Due to the low expression of GCN5a, we used four T-125 flasks for each biological replicate, with three biological replicates per sample.

Whole cell lysates were prepared by incubating the parasites in lysis buffer (50 mM Tris-HCl pH 7.4, 150 mM NaCl, 1 mM MgCl₂, 0.5% NP-40, 10% glycerol). After clarifying the lysates via centrifugation at 21,000 xg at 4°C, the lysate was precleared by incubating with mouse IgG magnetic beads (Cell Signaling, 5873S) for 4 h at 4°C with rotation. The supernatant was transferred onto α-HA magnetic beads (Thermo Fisher, 88836), which were pre-washed with lysis buffer, and incubated overnight with rotation at 4°C. Beads were washed with wash buffer (50 mM Tris-HCl pH 7.4, 150 mM NaCl, 1 mM MgCl₂, 0.05% NP-40, 10% glycerol) and with PBS and submitted for MS at the Proteomics Core Facility at IUSM as described before (44). Data were filtered to include only proteins that were not detected in the parental strains and were identified by at least 5 peptides in two out of three replicates.

### CUT&Tag

CUT&Tag was performed as described previously (45). Briefly, GCN5a^HA^, GCN5b^HA^, and untagged PruΔKu80ΔHX parental parasites were grown in confluent HFF monolayers for 48 h in tachyzoite or bradyzoite conditions. Parasites were isolated by syringe passage followed by filtration. Samples were processed using the CUT&Tag-IT assay kit (Active Motif, 53160) according to the manufacturer’s instructions with a primary antibody directed against the HA tag (Invitrogen MA5-27915). Sequencing reads were aligned to the *Toxoplasma* genome (v67) using bowtie2 allowing for dovetailing (46). Chromosome occupancy was visualized using the IGV genome browser (47). Data pertaining to this experiment can be obtained at GSE286088 (reviewer token: cvizwioknfepbmv).

### Chromatin immunoprecipitation (ChIP)

GCN5a^HA^ and untagged parental Pru parasites were cultured under alkaline stress for 48 h. Three independent biological samples were processed simultaneously for ChIP analysis. Parasites were fixed with 4% paraformaldehyde (Sigma, P6148) for 15 min, followed by PBS washes. Lysates were prepared as above for immunoprecipitation. Ten percent of the lysate was reserved for input. The remaining lysate was incubated onto α-HA magnetic beads and washed as above for immunoprecipitation. DNA was extracted from ChIP and input samples using phenol-chloroform (Invitrogen, 15593031) and precipitated with ethanol. qPCR was performed in duplicate, with Ct values normalized to input. Subtelomere-specific forward and telomere-specific reverse primers were used for analysis (Table S1).

### Fluorescence *in situ* hybridization of telomeres

Parasites were inoculated onto confluent HFFs and grown under bradyzoite-inducing conditions for 3 days. Monolayers were fixed in 4% paraformaldehyde for 15 min and incubated overnight at 4°C in 70% EtOH. Coverslips were processed as described in (48). Briefly, samples were denatured in 0.1N HCl for 15 min, then neutralized in 2X saline sodium citrate (SSC) (Corning, 46-020-CM) for 5 min before being placed in equilibration buffer (2X SSC, 50% formamide) for 30 min. Coverslips were incubated in hybridization buffer (2X SSC, 50% formamide, 15% dextran sulfate, 1% Tween-20) containing 2 μM of a telomeric probe (TTTAGGG_X4_-Cy5) overnight at 37°C. Coverslips were sequentially rinsed in 2X SSC, 1X SSC, 0.1X SSC, and PBS for 5 minutes then processed for IFA using anti-HA as above.

## ACKNOWLEDGEMENTS

This research was supported by grants from the National Institutes of Health (AI152583 and AI167662 to WJS). The authors thank Drs. Vern Carruthers (University of Michigan) and David Sibley (Washington University) for generously supplying antibodies used in this study. The mass spectrometry work performed in this work was done by the Indiana University School of Medicine Proteomics Core. Acquisition of the IUSM Proteomics core instrumentation used for this project was provided by the Indiana University Precision Health Initiative. The proteomics work was supported, in part, by the Indiana Clinical and Translational Sciences Institute (funded in part by Award Number UL1TR002529 from the NIH, National Center for Advancing Translational Sciences, Clinical and Translational Sciences Award) and, in part, by the IU Simon Comprehensive Cancer Center Support Grant (Award Number P30CA082709 from the NCI).

