## Supplemental Figures for "GCN5a is a telomeric lysine acetyltransferase whose loss primes *Toxoplasma gondii* for latency"

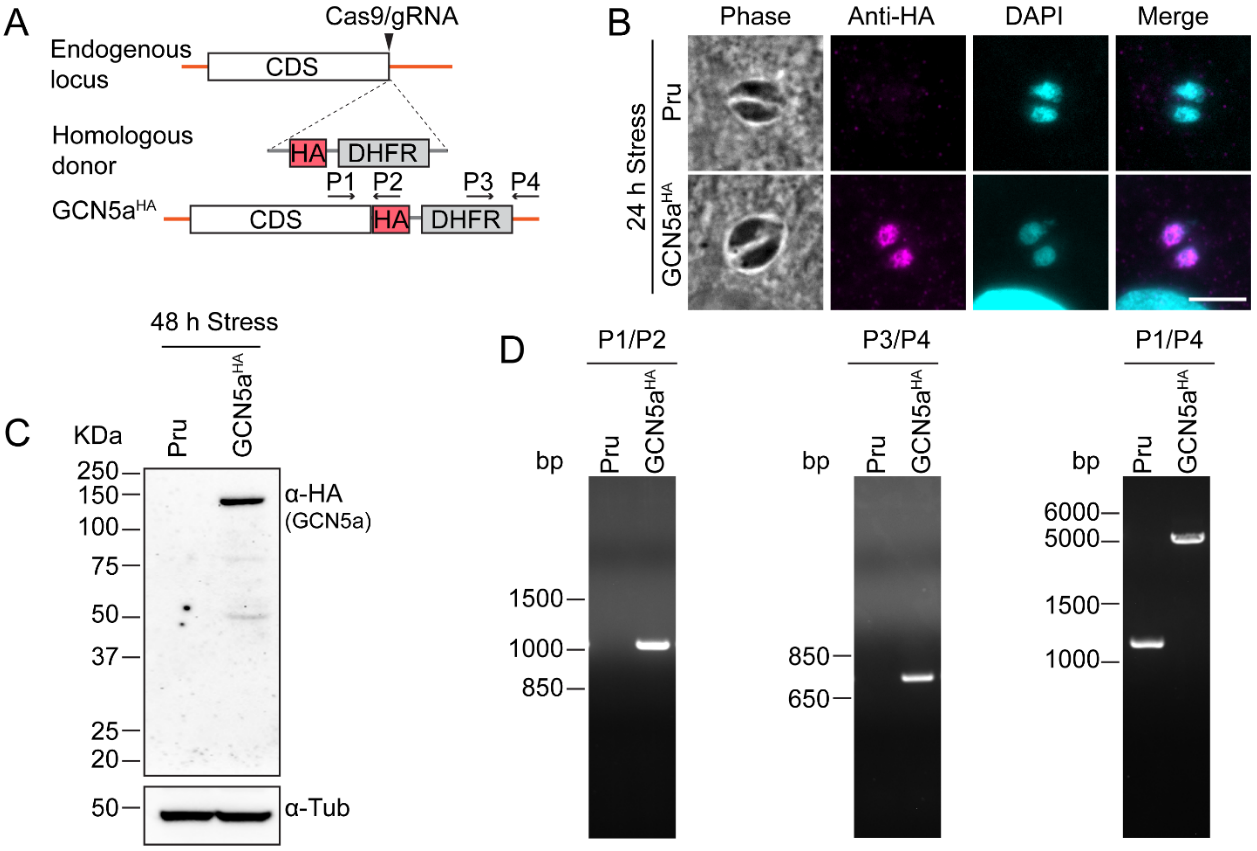


**Supplementary Figure 1. Endogenous tagging of GCN5a$^{HA}$.** A. Schematic of the *GCN5a* genomic locus and constructs used to tag the endogenous protein with an HA epitope at the C-terminus. B. IFA of Pru (parental) and GCN5a$^{HA}$ parasites following 24 h of alkaline stress. Anti-HA (magenta); DAPI (blue). Scale bar = 5 μm. C. Western blot of lysates made from Pru (parental) and GCN5a$^{HA}$ parasites after 48 h in alkaline stress probed with anti-HA. Tubulin served as a loading control. D. PCR analysis confirming the integration HA-DHFR using primers corresponding to the schematic in (A).

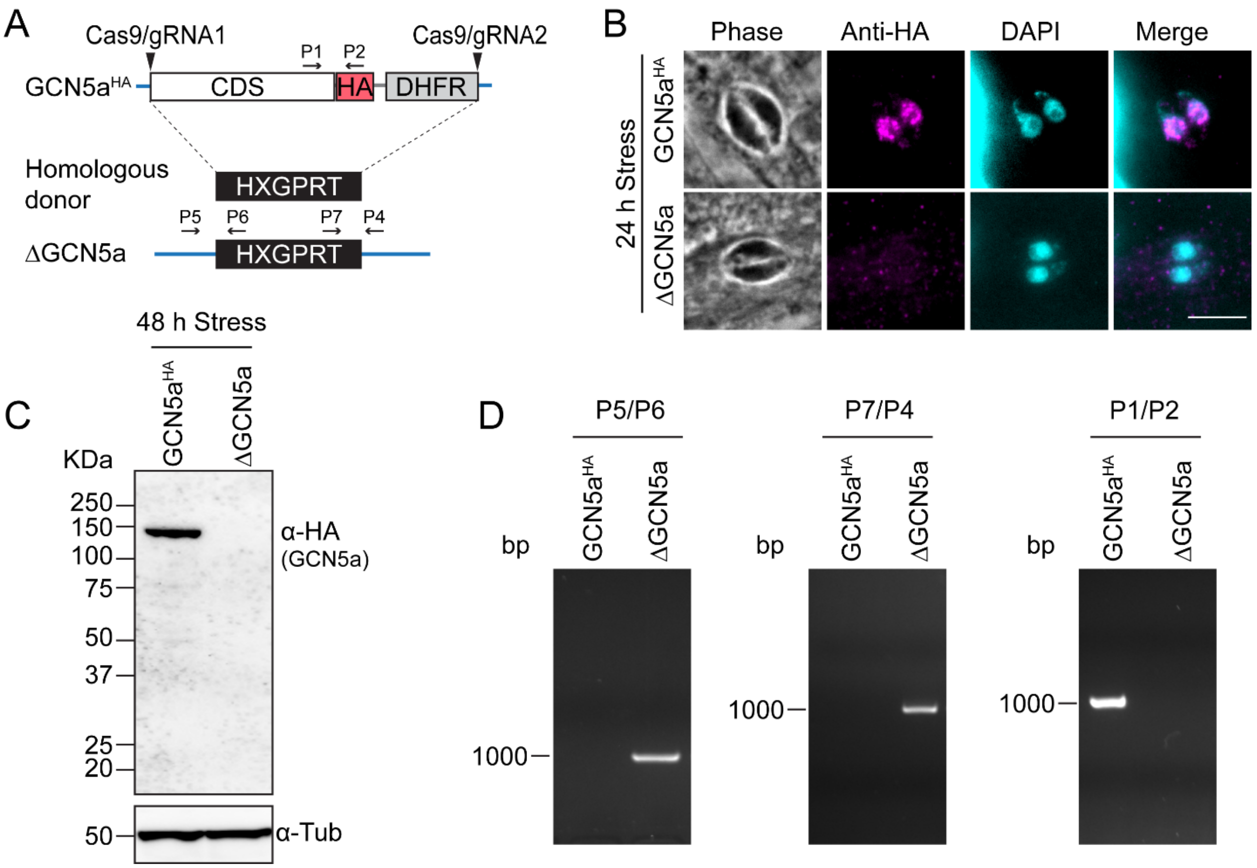


**Supplementary Figure 2. Generation of ΔGCN5a parasites.** A. Schematic of GCN5a$^{HA}$ genomic locus and HXGPRT construct to allelically replace it. B. IFA of GCN5a$^{HA}$ and ΔGCN5a parasites following 24 h of alkaline stress. Anti-HA (magenta); DAPI (blue). Scale bar = 5 µm. C. Western blot of lysates made from GCN5a$^{HA}$ and ΔGCN5a parasites after 48 h in alkaline stress probed with anti-HA. Tubulin served as a loading control. D. PCR analysis confirming the replacement of GCN5a$^{HA}$ with HXGPRT cassette using primers corresponding to the schematic in (A).

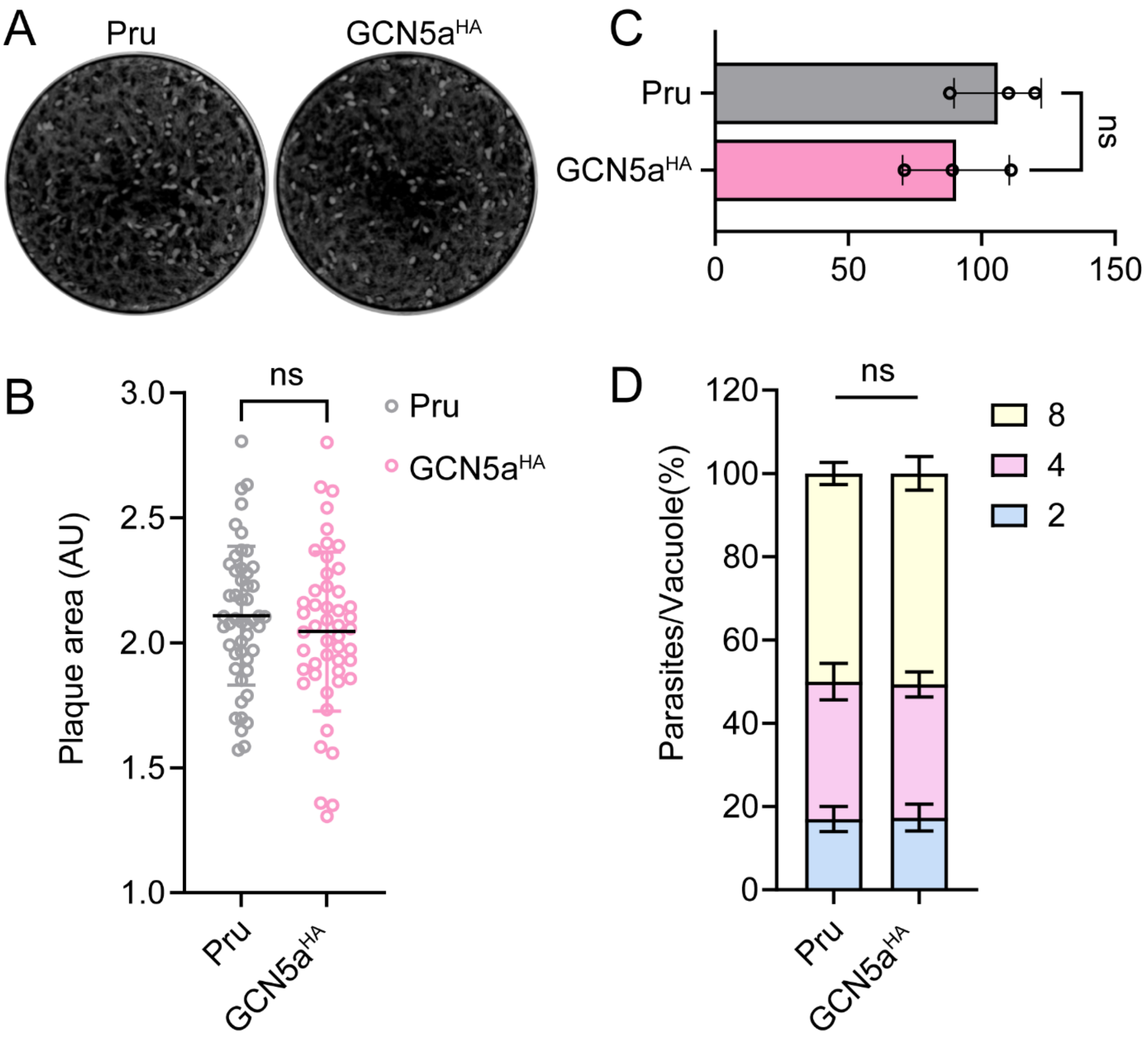


**Supplementary Figure 3. Growth assays comparing GCN5a$^{HA}$ and parental Pru parasites**. A. Plaque assay of Pru (parental) and GCN5a$^{HA}$ parasites grown under tachyzoite conditions for 12 days. B. The area of 50 randomly selected plaques was measured from 3 biological replicates. Statistical significance was measured by unpaired Student's t test; ns = not significant. C. Number of plaques formed on HFF monolayer by Pru and GCN5a$^{HA}$ parasites. Each data point represents an average of 3 biological replicates ± standard deviation. Statistical significance was measured by unpaired Student's t test; ns = not significant. D. Number of parasites per vacuole was counted from 100 randomly selected vacuoles. Statistical significance was measured by one-way ANOVA with Tukey's method for multiple comparisons; ns = not significant.

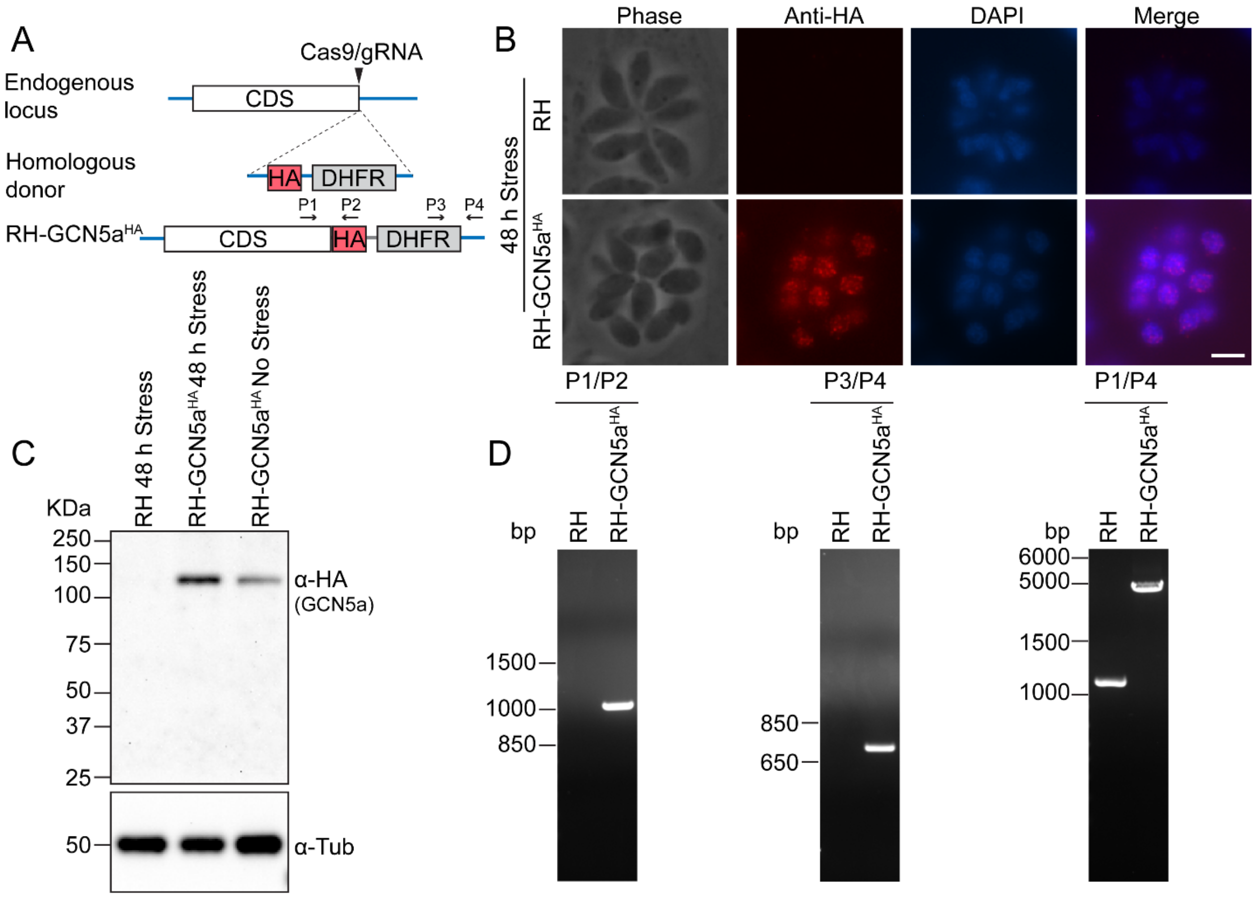


**Supplementary Figure 4. Endogenous tagging of GCN5a$^{HA}$ in RH strain.** A. Schematic of the *GCN5a* genomic locus and constructs used to tag the endogenous protein with an HA epitope at the C-terminus. B. IFA of RH (parental) and RH-GCN5a$^{HA}$ parasites following 48 h of alkaline stress. Anti-HA (red); DAPI (blue). Scale bar = 5 µm. C. Western blot of lysates made from RH (parental) and RH-GCN5a$^{HA}$ parasites after 48 h in alkaline stress probed with anti-HA. Tubulin served as a loading control. D. PCR analysis confirming the integration HA-DHFR using primers corresponding to the schematic in (A).

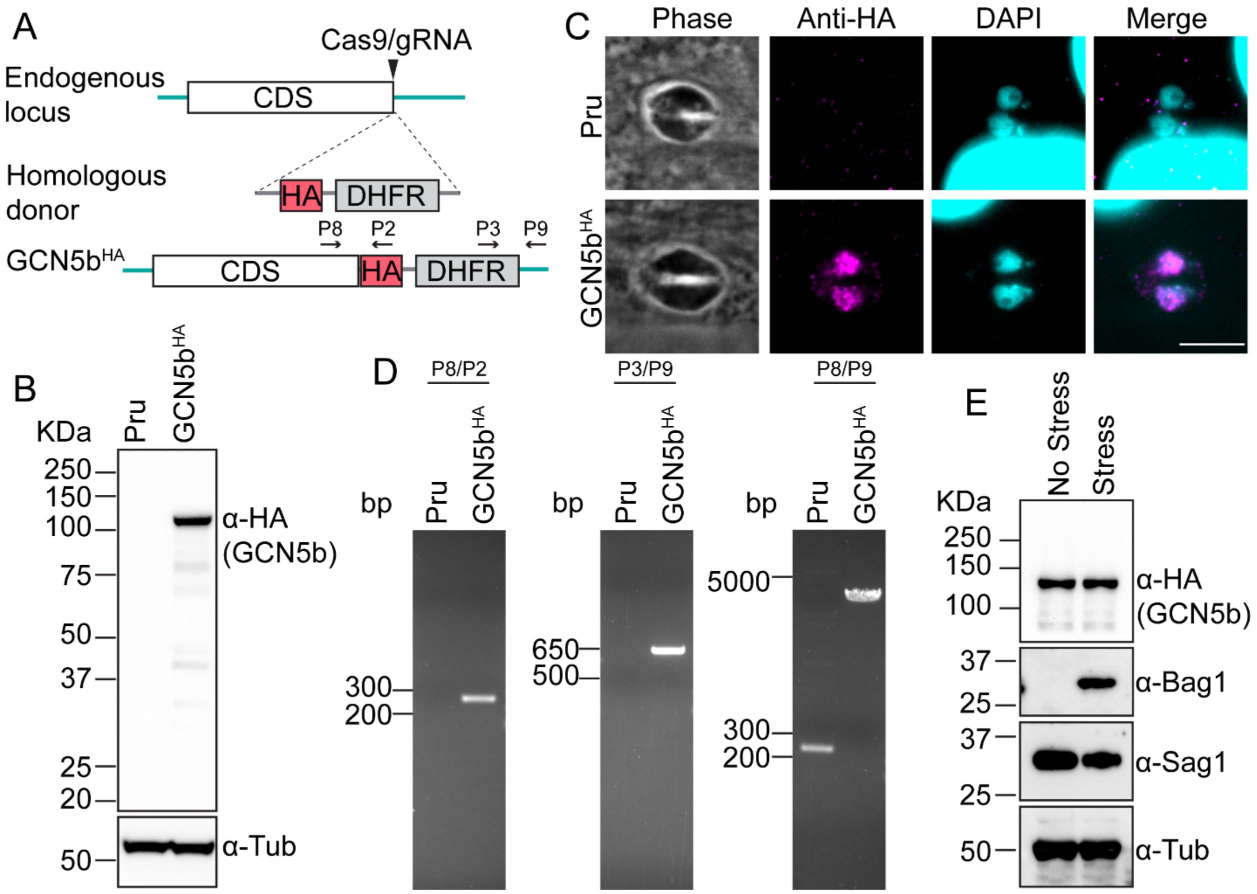


**Supplementary Figure 5. Generation and characterization of Pru GCN5b$^{HA}$ parasites.** A. Schematic of GCN5b genomic locus and constructs used to generate GCN5b protein endogenously tagged with an HA epitope at the C-terminus. B. Western blot of lysates made from Pru (parental) and GCN5b$^{HA}$ parasites probed with anti-HA. Tubulin served as a loading control. C. IFA of Pru (parental) and GCN5b$^{HA}$ parasites following 24 h of alkaline stress. Anti-HA (magenta); DAPI (blue). Scale bar = 5 µm. D. PCR analysis confirming the integration of HA-DHFR using primers corresponding to the schematic in (A). E. Western blot of GCN5b$^{HA}$ lysates grown under tachyzoite (No Stress) and alkaline stress (Stress) for 48 h probed with anti-HA and stage-specific markers BAG1 and SAG1. Tubulin served as a loading control.

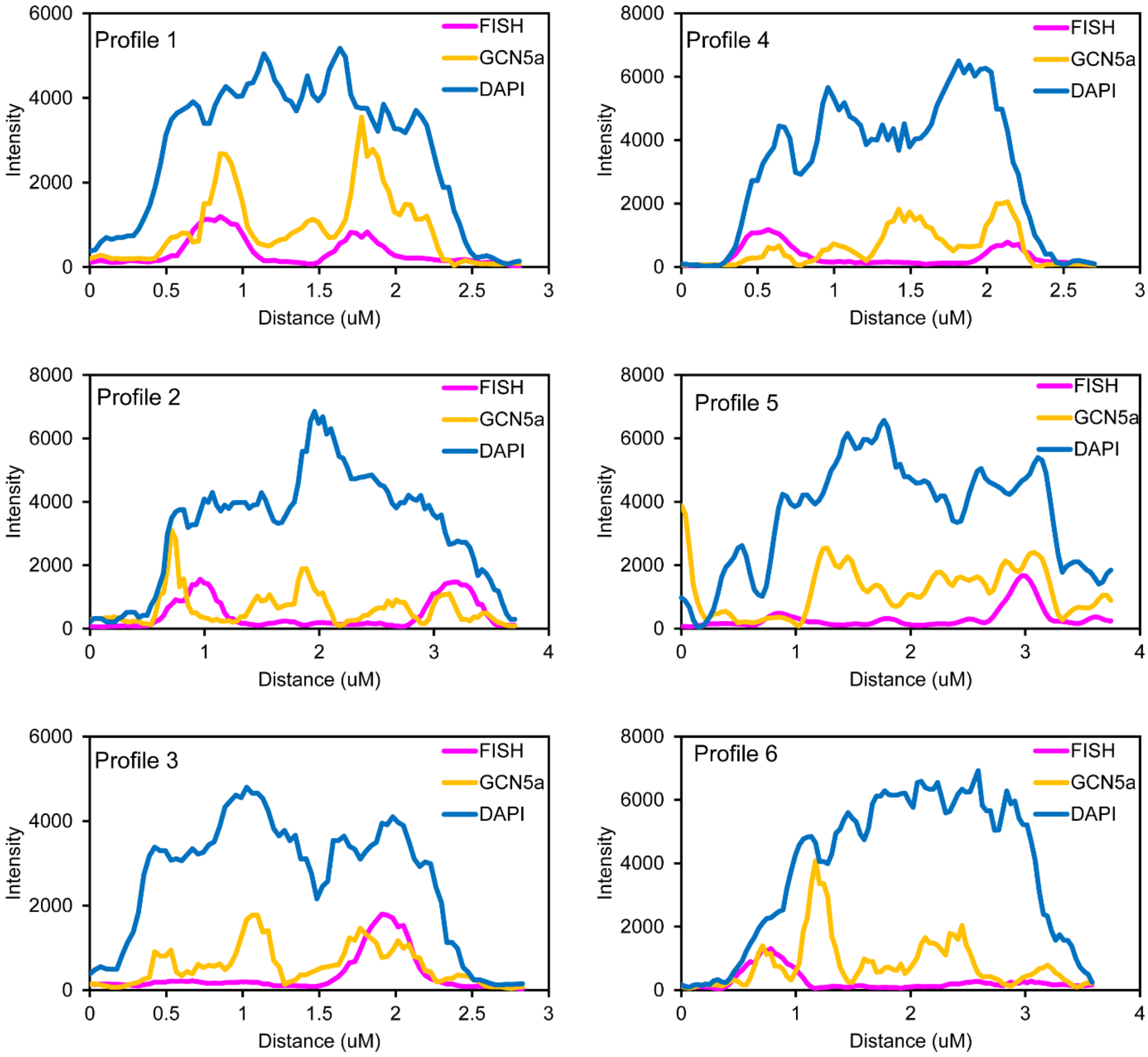


**Supplementary Figure 6. Additional densitometry profiles of GCN5a and telomere fluorescent signals.** Six profiles from the IFA image designated in Figure 6A were selected for analysis. The fluorescence intensity of each profile was measured and subsequently plotted to demonstrate partial localization of GCN5a to the telomeres.

### SUPPLEMENTAL TABLES

**Supplemental Table S1. List of oligonucleotides used in this study.**

**Supplemental Table S2. Differential expression analysis from RNAseq.**
